# Evidence for transposable element control by Argonautes in a parasitic flatworm lacking the piRNA pathway

**DOI:** 10.1101/670372

**Authors:** Anna V. Protasio, Kate A. Rawlinson, Eric A. Miska, Matt Berriman, Gabriel Rinaldi

## Abstract

Transposable elements (TEs) are mobile parts of the genome that can jump or self-replicate, posing a threat to the stability and integrity of the host genome. TEs are prevented from causing damage to the host genome by defense mechanisms such as nuclear silencing, where TE transcripts are targeted for degradation in an RNAi-like fashion. These pathways are well characterised in model organisms but very little is known about them in other species. Parasitic flatworms present an opportunity to investigate evolutionary novelties in TE control because they lack canonical pathways identified in model organisms (such as the piRNA pathways) but have conserved central players such as Dicer and Ago (argonaute) enzymes. Notably, parasitic flatworm Ago proteins are phylogenetically distinct from classical Ago, raising the question of whether they play special roles in these organisms. In this report, we investigate the role of Ago proteins in the parasitic flatworm *Schistosoma mansoni*. We show that transcript abundance of two retrotransposable elements increases upon silencing of *S. mansoni* Ago genes. We further demonstrate that SmAgo2 protein is primarily localised in the germ line of adult worms and its sub-cellular localisation is both nuclear and cytoplasmic. These findings provide further evidence of active TE control under a yet not fully unveiled pathway.

## Introduction

Genome integrity is an essential aspect of the biology for all organisms. Active mobile genetic elements, also termed transposable elements (TEs), pose a threat to this integrity as they can insert themselves randomly in the host genome [1]. Organisms from plants to humans have developed elegant molecular mechanisms to regulate TE activity. One such mechanism is nuclear silencing, in which transcripts arising from active TE genes are intercepted and degraded by ‘host’ mechanisms, ultimately preventing their insertion in the ‘host’ genome. At the centre of these pathways are Argonaute proteins. These are characterised by the presence of a PAZ domain and can be divided into two subfamilies: PIWI proteins and Ago subfamily. In animals, piRNAs (small 20-32 nucleotides RNA) associate with PIWI proteins and are at the centre of TE control in the male germ line [2]. In somatic tissues, Ago proteins associate with miRNA and other small RNAs and can participate in RNA induced silencing pathways both in the nucleus or the cytoplasm [3]. Notably, the mechanisms of nuclear silencing vary greatly across the tree of life, indicating independent evolutionary origins [2, 4].

Parasitic flatworms are a group of pathogens of clinical and veterinary relevance that cause some of the most prevalent and devastating infectious diseases affecting the poorest communities in the world [5, 6]. Ongoing work in the field of genomics has provided much insight into the genetic composition of these parasites. TEs constitute a sizable proportion of their genome: approximately 50% of the genome of all sequenced schistosome species, *Schistosoma haematobium* [7], *S. japonicum* [8] and *S. mansoni* [9] is comprised of repetitive elements.

In *S. mansoni*, TEs account for almost 40% of the genome and some are transcriptionally active [10, 11]. Despite this fact, very little is known about whether or how these parasites actively regulate TEs. Earlier studies revealed that schistosomes lack PIWI proteins [12] while the presence or absence of targeted DNA methylation remains controversial [13, 14, 15, 16]. Bioinformatics and phylogenetic analyses of schistosome genomes [12, 17] indicate that these parasites have three Ago genes, namely SmAgo1 (Smp_198380), SmAgo2 (Smp_179320) and SmAgo3 (Smp_102690)^1^. Phylogenetic studies revealed that SmAgo1 shares homology to Ago proteins of other organisms where the primary function is to mediate the canonical RNA interference pathway in the cytoplasm. On the other hand, SmAgo2 and SmAgo3 belong to a cluster of **FL**atworm-specific **A**go proteins (also called **FLA**go) [17]. The absence of PIWI proteins in parasitic flatworms has raised the question of how these worms regulate TEs [12].

Here we investigate a possible role for SmAgo proteins in TE regulation. We present evidence that silencing of SmAgo genes increases abundance of at least two retrotransposable elements in *S. mansoni*. Furthermore, we show that SmAgo2 protein can be localised to the worm’s reproductive system and in particuar, to the mature oocytes in the germ line of female worms, raising questions about a possible role in worm reproduction.

## Results

### Gene silencing of argonaute transcripts in *S. mansoni* increases abundance of retrotransposon transcripts

We employed dsRNA as a method for silencing transcript of Ago genes in *S. mansoni*. We generated dsRNA of approximately 700-800 bp for each of the three SmAgo transcript plus an irrelevant control (firefly luciferase) and a positive control against SmDNMT2. Adult worms were transfected with 30ug of dsRNA and kept in culture for 4 days. RT-qPCR results confirmed the reduction of all targeted transcripts (SmAgo1, SmAgo2 and SmAgo3, **Figure 1A**) as well as the positive control (SmDNMT2, data not shown). With this approach, we successfully silenced SmAgo1, SmAgo2 and SmAgo3 transcripts to 53%, 28% and 62% respectively (mean of three biological replicates).

**Figure 1.**
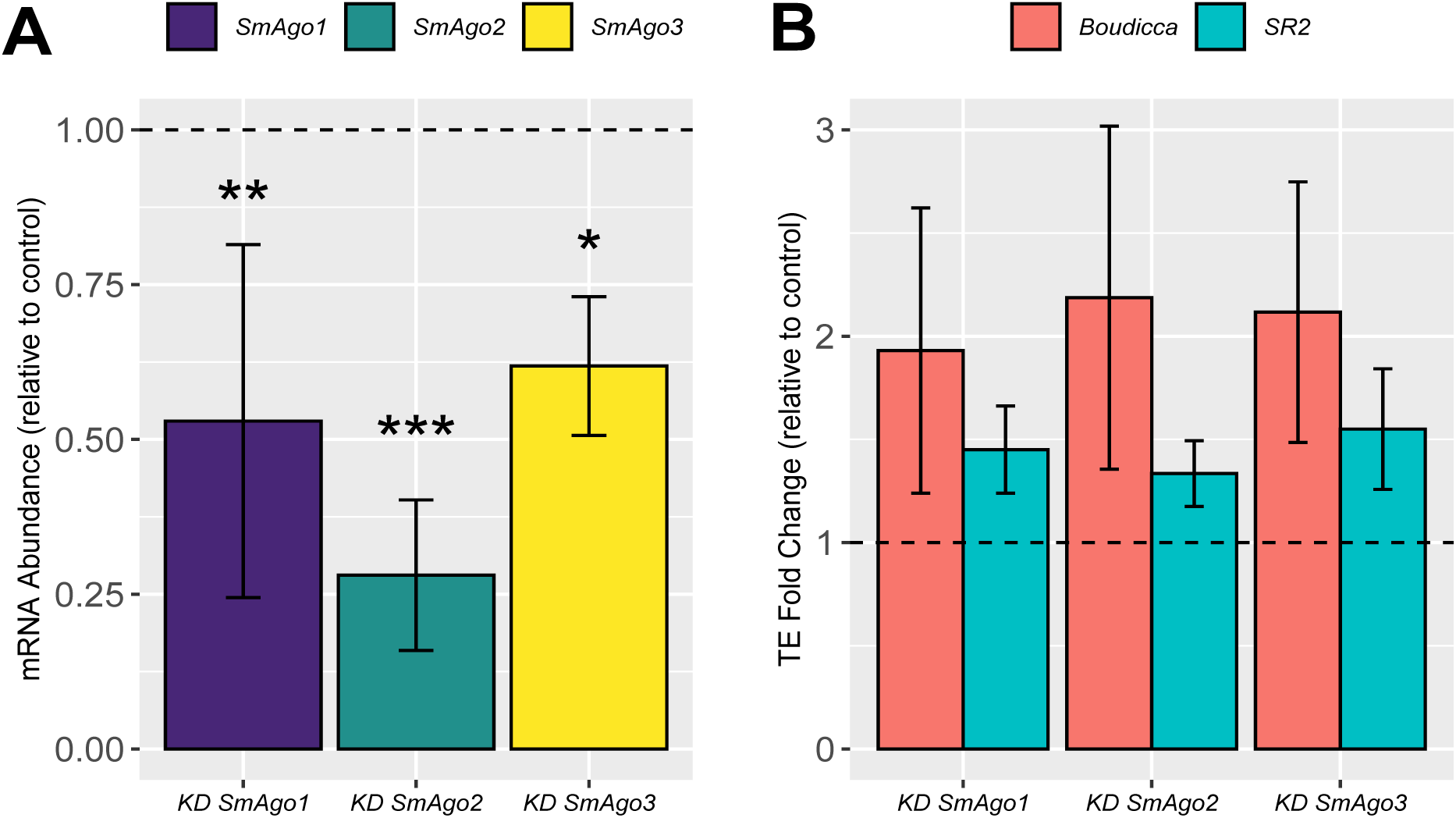
Quantification of Ago, Boudicca and SR2 transcripts in *S.mansoni* worms maintained *in vitro*. A) Reduced expression of SmAgo1, SmAgo2 and SmAgo3 in *in vitro* cultured worms. Barplot representing the relative abundance of target transcripts in dsRNA-treated samples. The horizontal dashed line at 1 represents the baseline of expression of target genes in the control sample. Bars represent the mean of three biological replicates, error bars represent ± 1 SE. (***) p-value < 0.001; (**) p-value < 0.01; (*) p-value < 0.05. B) Increase expression of retroelements Boudicca and SR2. Barplot represents the relative abundance of Boudicca and SR2 transcripts in samples of worms where SmAgo1, SmAgo2 or SmAgo3 were silenced. Dashed line set at 1 represents the baseline of expression of Boudicca and SR2 in control samples. Bars represent the mean of three biological replicates, error bars represent ± 1 SE.

Having demonstrated the successful silencing of SmAgo transcripts in our system, we wanted to investigate whether the reduction in their expression had an effect on the abundance of retrotransposon transcripts. To this end, we chose to focus on two well described *S. mansoni* retrotransposable elements, Boudicca and SR2. We measured transcript abundance of these two retroelements in SmAgo knock-downs and compared them to controls. We observed increased expression of Boudicca and SR2 in all SmAgo knock-down samples (**Figure 1B**). On average, Boudicca transcripts were twice as abundant in knock-down samples compared to controls, while in SR2, abundance increased but was less pronounced.

### SmAgo2 protein is localised in the nucleus and cytoplasm in adult schistosomes

Using a validated monoclonal antibody against *S. japonicum* SjAgo2 protein [18], we carried out immunofluorescence localisation of SmAgo2 protein in whole mounts as well as cross sections of *S*. *mansoni* adult worms (**Figure 2**). SjAgo2 shares 85% identity with SmAgo2 and the monoclonal antibody was generated against a peptide in the protein N-terminus where sequence identity with the other two SmAgo proteins is lowest. Our immunofluorescence results show that SmAgo2 protein localises to the ventral sucker and testes of the male and posterior ovary of the female. (**Figure 2, B** and **E**). No staining was observed in the negative control (omitting the primary antibody, Supplementary Figure 1).

**Figure 2.**
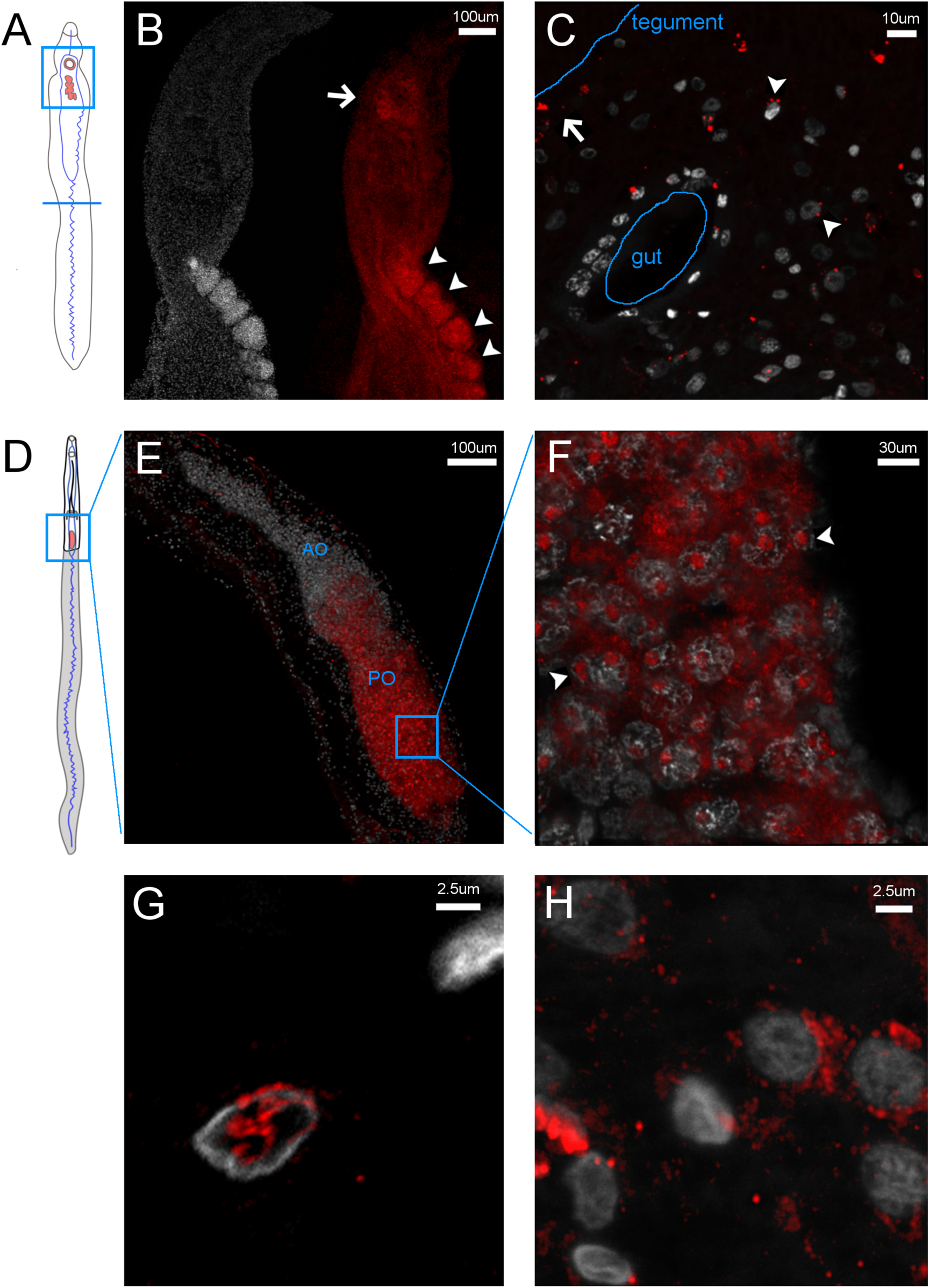
Immunofluorescence protein localisation of *S. mansoni* Argonaute2 (SmAgo2) in adult worms. In all panels, DAPI staining is represented in grey, SjAgo2 in red. A) Schematic of mature male worm showing overview of SmAgo2 localisation (pink-shaded areas); B) Whole mount of adult male reveals SmAgo2 localised in the testes (arrowheads) and ventral sucker (arrow); C) Cross section of male shows discreet distribution of SmAgo2 surrounding cell nuclei and possibly P-bodies (arrowheads) and also in close proximity to the tegument (arrow) where fewer cell nuclei are found; D) Schematic of mature female worm showing overview of SmAgo2 localisation (pink-shaded areas); E) Three-dimensional reconstruction of confocal image of the adult female ovary, PO posterior ovary, AO anterior ovary; F) Higher magnification maximum projection of confocal z-stack showing SmAgo2 in the cytoplasm and possibly in the nucleolus of mature oocytes; G) High magnification of cross section showing SmAgo2 localisation within a nucleus; and H) High magnification of cross section showing SmAgo2 localisation and in the cytoplasm of cells that sit within the parenchyma.

We were able to further characterise the localisation of this protein at the sub-cellular level. In a cross section of the male worm’s somatic tissue (**Figure 2C**), we observed that SmAgo2 is either diffuse in the cytoplasm or has a very discreet punctuated pattern similar to that shown by processing bodies [19]. Also termed P-bodies, these are dynamic centres for mRNA turnover, micro RNA activity and mRNA silencing. Argonaute proteins in other species have also been observed in P-bodies as well as diffuse in the cytoplasm [19, 20]. We also detected SmAgo2 immediately under the tegument, in a region with fewer cell nuclei. SmAgo2’s proximity to the constantly shedding adult tegumental surface suggests that it could be shed alongside the tegument into the bloodstream of the mammalian host.

We detect SmAgo2 in the mature ova in the posterior ovary (**Figure 2E**). In contrast, no SmAgo2 is observed in less developed primary or secondary oocytes (immature ova) found in the anterior ovary, where active cell division occurs [21, 22]. Mature ova are characterised by the large size of the nucleus, the well defined nucleolus and scattered chromatin fibres [21]. These were clearly distinguishable in our DAPI staining (Supplementary Figure 2). Within these cells, SmAgo2 localisation resembles that of a nucleolus (**Figure 2F**), but additional co-localisation experiments will be required to verify this. SmAgo2 is also found in the very thin layer of mature oocyte cytoplasm that surrounds the oocyte nucleus (**Figure 2F**). Whereas SmAgo2 seems to be located both in the nucleus and cytoplasm in mature oocytes, in somatic cells it locates to either one or the other space but not both (**Figure 2, E** and **F**).

## Discussion

Control of TEs is an essential cell function of all organisms. Parasitic flatworms lack classical TE control pathways mediated by the PIWI proteins and their associated piRNAs or the RNA-dependent RNA-polymerase [12]. In this work, we used experimental approaches to further characterise the FLAgo enzymes of *S. mansoni*. We chose *S. mansoni* because of its relatively simple life cycle maintenance but we believe that these important molecular pathways are likely to be shared among other members of the genera. We successfully achieved significant reduction of transcripts from the three *S. mansoni* Ago genes and we further demonstrated that such silencing is associated with increased abundance of two retrotransposable elements, namely Boudicca and SR2. Boudicca was the first full-length LTR retrotransposon described in *S. mansoni* [11], while SR2 is the non-LTR retrotransposon most closely related to the RTE-1 family of non-LTR retrotransposons found in *C. elegans* [23]. They have both been reported to have between 1,000 and 10,000 copies in the genome and they are readily detected in RT-qPCR experiments. Although increased abundance of Boudicca and SR2 transcripts was consistently recorded in our SmAgo knock-down samples, the variance for each replicate group was very high probably reflecting variable penetrance of the phenotype in each experiment and perhaps in each of the 8-10 individual worms present in each replicate.

The results obtained for SmAgo1 knock-downs were at first puzzling, given that sequence similarity and phylogeny suggest that this protein is involved in cytoplasmic RNAi. Flatworms have a reduced complement of argonaute proteins in comparison to other organisms such as *C. elegans*, Drosophila, mouse and human [17], and they lack PIWI proteins [12]. Therefore, it is possible that functional redundancy exists among the relatively small complement of flatworm Ago proteins, allowing them to fulfil the same roles taken by more specialised Argonaute proteins in other organisms.

Continuing with our characterisation of SmAgo function, we sought to investigate their tissue localisation. We found SmAgo2 protein in the reproductive system of both male and female adult worms as well as in the main male attachment organ, namely the ventral sucker. These findings are in agreement with previous reports on the gene expression patterns of SmAgo2. Whole mount *in situ* hybridisation experiments showed SmAgo2 expression in adult worms in the posterior ovary and vitelline glands in the female and testes of male worms and attachment organs [24] as well as stem-cells [22]. The ovary of sexually active females is long and pear-shaped and is divided into anterior and posterior section. Oocytes originate in the anterior lobe and mature as their progress to the posterior ovary [21]. We detect SmAgo2 protein in the female posterior ovary where it is present exclusively in the mature oocytes. The very distinct sub-cellular distribution suggests possible nucleolar localisation. Argonaute proteins have been detected in the nucleolus of plants [25], human cells [26] and most recently in the fruit fly’s ovarian somatic cells, where evidence points to a role in suppressing expression of TEs inserted between ribosomal DNA copies [27].

A family of vasa-like genes (*Smvlg1-3*) identified in schistosomes have also been localised to the posterior ovary [28] and are essential in the development and maintenance of schistosome reproductive organs [29, 30]. Vasa genes are highly conserved RNA helicases essential for germ line development, probably through their role as assembly docks for the piRNA silencing complex, which includes an Argonaute protein, and whose ultimate goal is TE degradation [31]. Whether vasa-like proteins interact with SmAgo2 in *S. mansoni* to carry out similar roles remains to be investigated.

In the absence of PIWI proteins in trematodes and other parasitic flatworms, we speculate that the functions carried out by the many Argonaute proteins in other species (at least five in Drosophila and up to 25 in *C. elegans* [20, 3]) are concentrated in the relatively small number of parasitic flatworm Argonaute proteins, requiring them to be more functionally versatile.

## Materials and Methods

### Parasite material

The *S. mansoni* life cycle is maintained at the Wellcome Sanger Institute. Balb/c female mice were infected with 250 *S. mansoni* cercariae either intraperitoneally or percutaneously and adult worms were recovered six weeks post infection through portal perfusion. Parasites were washed extensively in DMEM and incubated in Adult Worm Media for 24 hours prior to either dsRNA treatment or fixation for immunofluorescence. Adult Worm Media: DMEM (5.4 g/L D-Glucose, Sigma) supplemented with Antibiotic Antimycotic (Thermo Fisher Scientific, UK) and 10% fetal calf serum.

### Double-stranded RNA (dsRNA) design and synthesis

Primers that incorporate the T7 promoter sequence at the 5’-end were designed to amplify a region of 500-800 bp of SmAgo2 and SmAgo3 (Supplementary Table 1). End-point PCR was performed using cDNA from *S. mansoni* adults as a template, 1 uM final concentration of a mix of SmAgo2 or SmAgo3 forward and reverse primers (Sigma, UK), and 2x Qiagen Fast Cycling PCR master mix (Qiagen, cat. no. 203743), following manufacturer’s instructions and 55*°*C annealing temperature. PCR products were checked for expected size and cloned into pCR-TOPO TA vector (Invitrogen, UK). Plasmid transformation and propagation was performed in heat-shock competent BL21 *Escherichia coli* cells [32]. A plasmid clone containing a section of SmAgo1 inserted between two T7 promoters was kindly donated by Dr James Collins (UT Southwestern, Dallas, U.S.A.). Plasmids were then isolated using QIAprep Spin Miniprep (Qiagen, cat no 27104), sequenced and used as templates to generate PCR products. These were then used as templates for *in vitro* transcription using a MEGAscript T7 Transcription Kit (Invitrogen, cat no AM1334) according to manufacturer’s instructions. A dsRNA integrity check and quantification was performed in agarose gels plus ethidium bromide staining against a ladder of known concentration. The dsRNA stocks were kept at −80*°*C in nuclease-free water at a concentration of 1 ug/ul.

### Parasite transfection with dsRNA

Five to eight *S. mansoni* adult worms were transfected with 30 ug of dsRNA targeting either SmAgo1 (Smp_198380), SmAgo2 (Smp_179320), SmAgo3 (Smp_102690), the irrelevant control firefly luciferase or positive control SmDNMT2 (Smp_334230)^2^. Parasites were placed in a 4 mm electroporation cuvette (BioRad, UK) containing 100 ul of simple media and siRNA/dsRNA and incubated for 10 mins in ice. Electroporation was performed with a single pulse at 125 V for 20 ms using a BTX Gemini X2 (BTX), immediately followed by addition of pre-warmed (37*°*C) full media [33]. Parasites were transferred to 6-well plates and incubated at 37*°*C and 5% CO_2_. Cultures were checked daily to control parasite viability and one-third of the culture media volume was replaced every other day. No significant loss of normal phenotype was detected (when compared to a non-electroporated control). On the day of parasite collection, worms were transferred to a 1.5 ml centrifuge tube and washed extensively with simple media. After the final wash, all media was removed, ml of Trizol reagent (Invitrogen, UK) were added to the tubes and these were kept at −80*°*C until RNA extraction. Simple media: DMEM (5.4 g/L D-Glucose, Sigma) supplemented with Antibiotic-Antimycotic (Thermo Fisher Scientific, UK). Full media: Simple media supplemented with 10% fetal calf serum and 2 mM L-glutamine.

Commercially sourced siRNAs designed against SmAgo1-3 were also used to attempt RNAi but we could not achieve reproducible silencing of these genes.

### RNA extraction and real-time quantitative PCR

RNA extraction was performed using a standard phenol-chloroform method (for full protocol see Trizol reagent user manual) where RNA was precipitated using 1 volume of isopropanol per volume of aqueous phase extracted. RNA was resuspended in nuclease-free water, assayed for integrity and quantification of nucleic acids in an Agilent Bioanalyzer. Poly-A enrichment prior to cDNA synthesis was performed using a Dynabeads mRNA Purification Kit (Thermo Fisher Scientific, UK, cat no 61006) according to manufacturer instructions. Superscript III Reverse Transcriptase (Invitrogen, UK, cat no 10368252) was used for cDNA synthesis. Real time quantitative PCR was performed with KAPA SYBR FAST qPCR Master Mix (Sigma-Aldrich - Merck, UK, cat no KK4602) according to manufacturer’s instructions in a StepOnePlus Real-Time PCR System (Thermo Fisher Scientific, Applied Biosystems) and using two endogenous controls: alpha-tubulin (Smp_090120) and GAPDH (Smp_056970).

Primer sequences are available in Supplementary Table 1.

### Statistical Analysis and visualisation

Two main changes in transcript abundance were analysed: i) change in expression of knocked-down genes (SmAgo1, SmAgo2 and SmAgo3 plus positive control SmDNMT2) and ii) change in expression of Boudicca and SR2. In all cases, the relative change abundance of a given transcript is normalised against the same transcript in the dsRNA irrelevant control (firefly luciferase, represented with black dashed lines in Figure 1). Therefore, all relative expression is calculated against the same gene but in the untreated sample. Student paired T-test implemented in R [34] was used to test the difference between sample means. Figure 1 was prepared using the ggplot2 package [35] implemented in R. Figure 2, Supplementary figures 1 and 2 were assembled using the free software GIMP [36].

### Immunofluorescence and histology

Adult worms were relaxed in 0.6 M of MgCl_2_ for 1 minute before fixation in 4% paraformaldehyde (PFA) in PBSTx (1 x PBS + 0.3% Triton-X 100) for 4 hours at room temperature. They were rinsed in PBSTx (3 × 5 mins, 2 × 1 hour washes) at room temperature and dehydrated in a step-wise ethanol series. For whole mount preparations, worms were rehydrated into 1xPBSTx, treated with 2mg/mL Proteinase K for 15 mins at 37*°*C, re-fixed in 4% PFA for 20mins, rinsed in PBSTx, blocked for 1 hour in 10% heat-inactivated sheep serum in PBSTx, and incubated overnight in primary antibody (see below) at 4*°*C. The worms were then rinsed in PBSTx (3×10mins, 3 × 1hr) and incubated in secondary antibody overnight at 4*°*C. Worms were rinsed in PBS (3×10mins, 3 × 1hr) and equilibriated in mounting media with 4’,6-diamidino-2-phenylindole DAPI (Fluoromount G, Southern Biotech, Birmingham, AL) overnight before mounting and imaging.

For paraffin sections, dehydrated worms were cleared in histosol (National Diagnostics) for 20 mins, and embedded in paraffin overnight. Paraffin blocks were sectioned at 8 um using a Leica (RM2125 RTF) microtome. For immunostaining of paraffin sections, slides were dewaxed in Histosol (2×5 min), then rehydrated through a descending ethanol series into PBS + 0.1% Triton (PBT, 2×5 min). Slides were blocked with 10% heat-inactivated sheep serum in PBT for 1 hour at room temperature in a humidified chamber. The primary antibody (see below) was diluted in block (10% heat-inactivated sheep serum in PBT) and applied to the slide, covered with parafilm, and incubated at 4*°*C for 48 hours. Slides were then rinsed in PBT (3×10min). A secondary antibody diluted in block solution were then applied to each slide, and slides were covered with parafilm and incubated in a humidified chamber, in the dark, at room temperature for 2 hours. Slides were rinsed in PBT 3 × 10 mins, and then 4 × 1 hour prior to counterstaining with the nuclear marker (DAPI) (1 ng/ml) and mounting in Fluoromount G (Southern Biotech, Birmingham, AL). The primary antibody used was anti-*Schistosoma japonicum* Ago2 (clone 650-1-1-KM864-3-11E8) from Abcam lnc (NJ, US) with kind permission from Dr Pengfei Cai (QIMR, Australia). This was used diluted 1:200. An anti-mouse secondary antibody was used at 1:500 dilution. A negative control was obtained by omitting the primary antibody and no fluorescence signal was detected. Imaging was carried out using an epi-fluorescent Zeiss Axioscope microscope.

## Supplementary Tables

**Supplementary Table 1.**
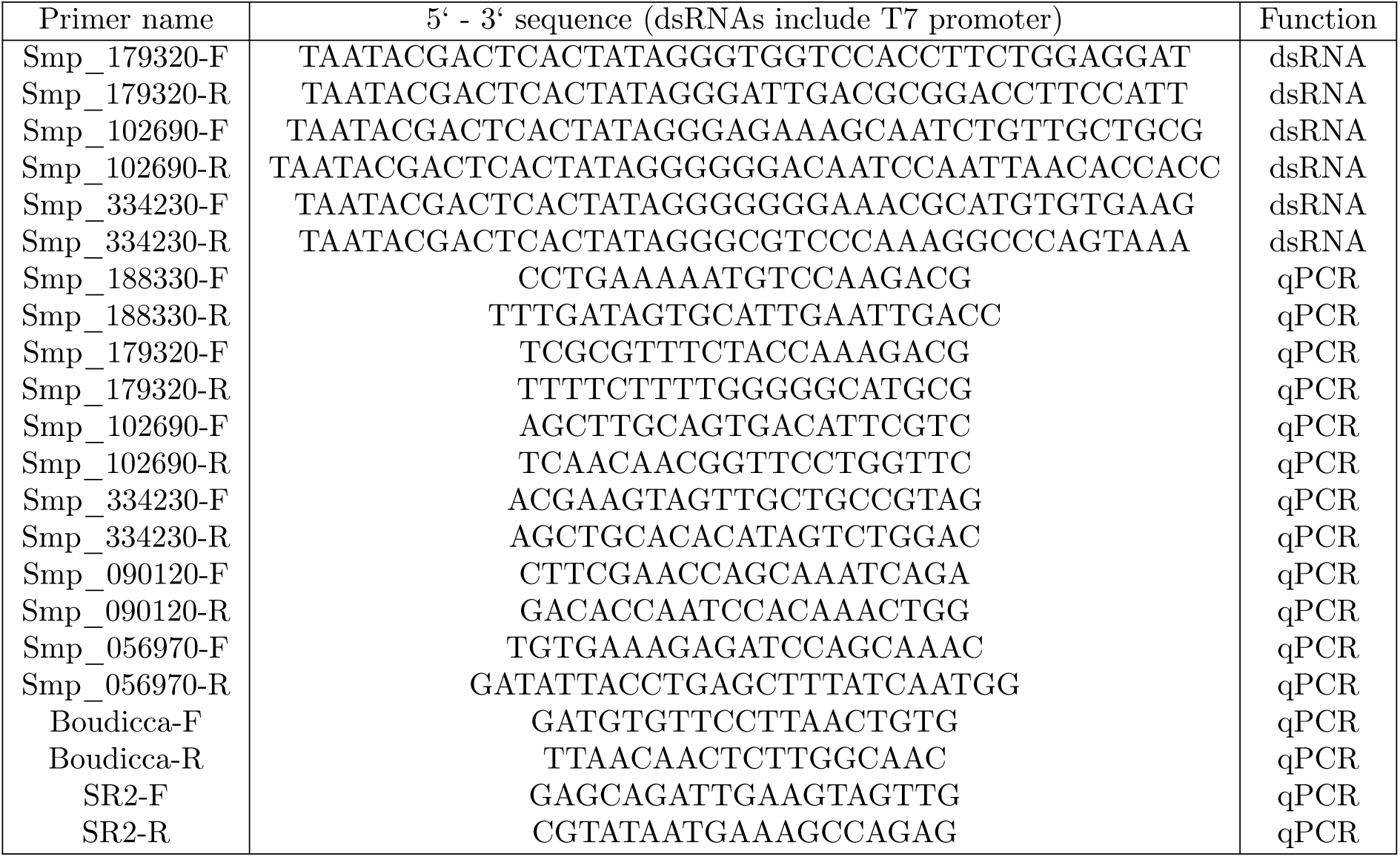
Primer sequences used for dsRNA generation and real-time PCR.

## Supplementary Figures

**Supplementary Figure 1.**
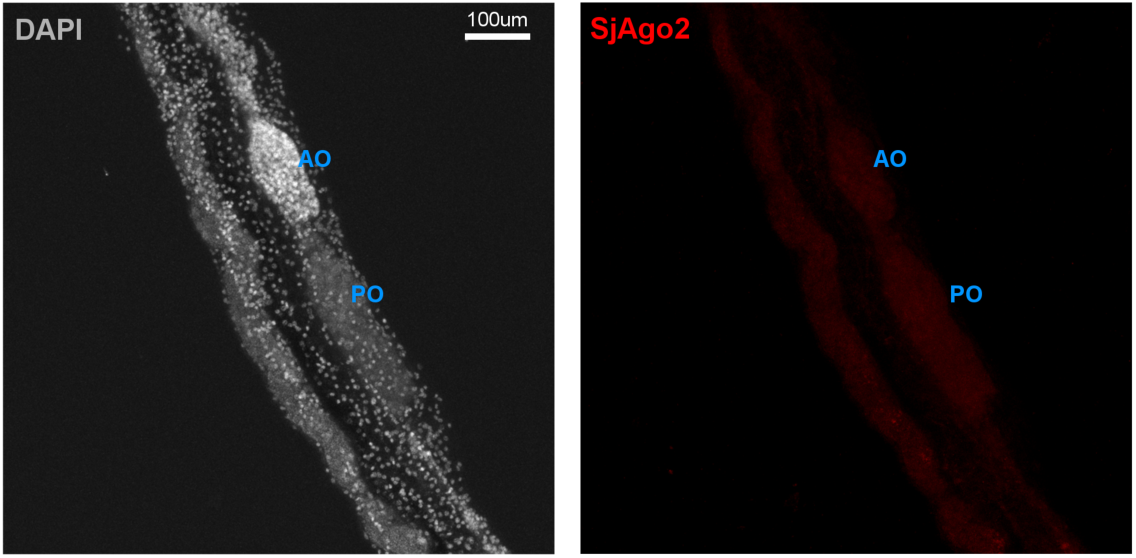
Immunofluoresence control experiment (with primary antibody omitted). Optical section of female whole mount showing anterior (AO) and posterior ovary (PO).

**Supplementary Figure 2.**
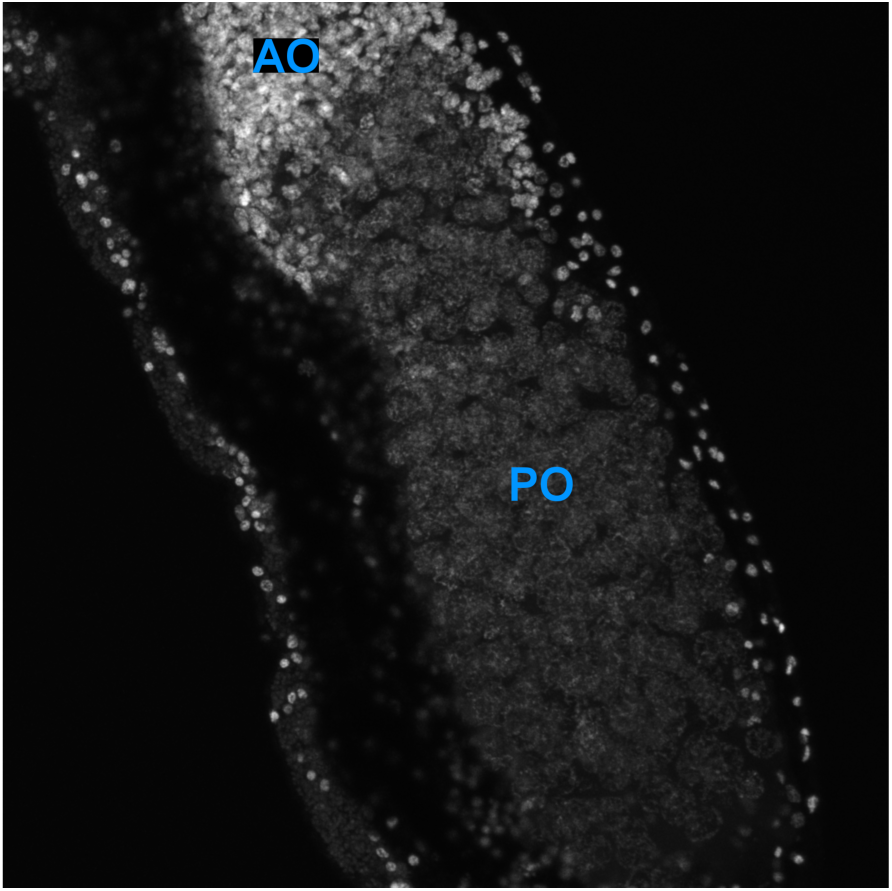
DAPI staining of the ovary of a female worm. Notice the normal size of the nuclei of immature oocytes in the anterior ovary (AO) and the enlarged size of nuclei of the mature oocystes in the posterior ovary (OV).

## Declarations

### Ethics Approval

Mouse infections to maintain the life cycle of NMRI strain of *S. mansoni* and collect adult worms were conducted under the Home Office Project Licence No. P77E8A062, and all protocols were approved by the Animal Welfare and Ethical Review Body (AWERB) of the Wellcome Sanger Institute. The AWERB is constituted as required by the UK Animals (Scientific Procedures) Act 1986 Amendment Regulations 2012.

### Consent for publication

Not applicable.

### Availability of data and materials

The materials and datasets used and/or analysed during the current study are available from the corresponding author on reasonable request.

### Competing interests

The authors declare that they have no competing interests.

## Funding

This work was funded by Cancer Research UK (C13474/A18583, C6946/A14492) and the Wellcome Trust (104640/Z/14/Z, 092096/Z/10/Z) to EAM, a Wellcome Trust Janet Thornton Fellowship (WT206194) to KAR, and Wellcome Snager Institute core-funding support (grant WT206194).

## Authors’ contributions

AVP - Conception and design of the work, data acquisition, analysis, and interpretation of data. Preparation of the manuscript. KAR - Data acquisition, analysis, and interpretation of data. EAM - Provided materials and reagents. MB - Provided materials, reagents and laboratory space. GR - Provided materials and reagents and contributed to data interpretation.

## Acknowledgements

We are very grateful to: Dr Pengfei Cai (QIMR, Australia) for granting us use of the SjAgo2 antibody, Dr James Collins (UT Southwestern, Dallas, U.S.A.) for kindly providing us with SmAgo1 plasmid. We also like to thank our colleagues at the Wellcome Sanger Institute: Dr Simon Clare, Cordelia Brandt, and Catherine McCarthy, for assistance and technical support with animal infections and maintenance of the *S. mansoni* life cycle; David Goulding and Claire Cormie of the Electron and Advanced Light Microscope Facility for their support and assistance with imaging.

## Footnotes

1 Smp_ names correspond to Wormbase Parasite accession numbers, https://parasite.wormbase.org/

2 Smp_ names correspond to Wormbase Parasite accession numbers, https://parasite.wormbase.org/

## Notes

### Competing Interest Statement

The authors have declared no competing interest.

### Summary of Updates

Syntax and supplementary tables fixed.

